# The effect of transcranial direct current stimulation over the dorsolateral prefrontal cortex on behavioral and brain activity indices of visuospatial attention in intrinsic reward contexts

**DOI:** 10.1101/2023.08.01.551490

**Authors:** Atakan M. Akil, Renáta Cserjési, Dezső Németh, Tamás Nagy, Zsolt Demetrovics, H. N. Alexander Logemann

## Abstract

Research indicates a connection between frontal brain activity asymmetry and self-regulation, particularly, approach and inhibitory tendencies. However, the underlying brain mechanism remains unclear. Our preregistered study used transcranial direct current stimulation (tDCS) to overcome limitations in prior correlational studies, investigating the link between frontal alpha asymmetry (FAA), a potential neuromarker and a treatment target for relevant psychiatric disorders, and the behavioral and brain activity components related to approach and avoidance tendencies, as observed in a visuospatial cueing (VSC) paradigm. We utilized a randomized sham-controlled design with 65 healthy participants. Participants’ resting-state EEG was recorded to calculate FAA scores before and after 2 mA anodal tDCS to the right frontal site. They also completed a VSC task with neutral and intrinsic reward-associated (food) conditions. Results indicated no impact of tDCS on FAA or behavioral attentional bias/disengagement. Surprisingly, secondary analyses revealed tDCS enhancing attentional bias for rewards, as seen in enhanced Late Directing Attention Positivity and P1 effect. However, these effects did not translate into observable behavioral changes. The observed effects are consistent with a noradrenergic mechanism rather than asymmetry of brain activity.

## 1. Introduction

Self-regulation has been defined as “the self’s ability to control its own thoughts, emotions, and actions” (Baumeister et al. 1994), and its failure has been associated with mental health problems, such as depressive disorders and addictive disorders (Johnstone et al. 2007; Heatherton and Wagner 2011). Previous studies also indicated that self-regulatory dysfunction is the main cause of numerous contemporary problems like obesity and sexual infidelity (Heatherton and Wagner 2011). One plausible neural correlate of self-regulation, and notably approach and avoidance components, is frontal alpha asymmetry (FAA), which may also serve as a treatment target for relevant psychiatric disorders. More specifically, FAA is considered to be driven (at least in part) by activity within the dorsolateral prefrontal cortex (DLPFC) and represents a relative measure of the difference in alpha power (8-13 Hz) between the left and right cortices (Smith et al. 2017). Since the amplitude within the alpha frequency band represents inactivity, lower frontal asymmetry scores (right minus left alpha) reflect relatively less left than right cortical activity (Harmon-Jones et al. 2010). This asymmetry in frontal cortical activity has been associated with the two antagonistic components of self-regulation: approach and avoidance (Amodio et al. 2008; M.J.F. Robinson, A.M. Fischer, A. Ahuja 2016). In particular, cues signaling reward trigger approach motivation, which is represented by left-over-right frontal brain activity, while cues signaling threat activate inhibitory motivation that is represented by right-over-left frontal brain activity (Coan et al. 2006; Harmon-Jones et al. 2010; Kelley et al. 2017). Hence, these mechanisms allow individuals to deal with rewards and threats that arise in their environment. Interestingly, assumed manipulation of FAA via transcranial direct current stimulation (tDCS) has been suggested to affect approach relative to avoidance tendencies (Kekic et al. 2017). However, it remains uncertain whether tDCS-induced effects on approach tendencies towards reward-associated stimuli are mediated by tDCS-induced changes in asymmetry of frontal brain activity. Furthermore, the specific components underlying approach tendencies and their potential susceptibility to tDCS effects have not been elucidated. The present study aimed to investigate and provide answers to these questions.

It should be noted that approach tendencies and inhibitory control can be captured in a Posner paradigm, or visuospatial cueing (VSC) task (Posner et al. 1980). To elaborate, in the realm of visuospatial attention, attentional bias that presumably drives approach tendencies, and disengagement of attention associated with inhibitory control, are two crucial components (Posner et al. 1980). When an observer anticipates a target at a particular position in a visual scene, a top-down mechanism resulting in an attentional bias is associated with an accelerated processing of the target at the attended location (Corbetta and Shulman 2002). Conversely, attentional disengagement is a target-driven bottom-up mechanism that has conceptual overlap with inhibitory control (Posner et al. 1980; Logan et al. 1984; Logemann et al. 2017). When the observer has to focus on a more important object at the unattended site, it simplifies the decoupling of attention from the attended location, which simply facilitates a reorientation of attention (Logemann et al. 2014a, 2014b).

The VSC Task is a computer task used to investigate how attention is oriented in response to visual cues, which indicate the potential location of targets, and targets themselves (Posner et al. 1980). The most important behavioral outcome is the difference in response times (RTs) for validly cued targets and invalidly cued targets, which is called “validity effect,” and it is thought to reflect both attentional bias and attentional disengagement (Clark et al. 1989).

Earlier research has suggested that the magnitude of a reward (e.g., stimuli with high or low rewarding value) impacts attentional disengagement. Specifically, stimuli associated with greater rewards tend to sustain attention, even when detrimental to ongoing task performance (Watson et al. 2020). Moreover, there has been discussion regarding the efficiency of the tasks utilized to measure the engagement and disengagement of attention (Clarke et al. 2011). However, such studies often intermix neutral stimuli, rewarding distractors, and target stimuli within the same blocks, making it challenging to disentangle the distinct attentional processes in various conditions. Besides, these studies primarily relied on eye-tracking.

In the current study, similar to previous studies, we segregated neutral and reward blocks to circumvent this issue (Tsegaye et al. 2022). In contrast to previous research that depended on learned rewards (e.g., money), we employed an intrinsic reward condition involving appetizing food pictures, drawing from an earlier study that highlighted differences between conditions in terms of inhibitory control (Tsegaye et al. 2022). Furthermore, we combined the VSC Task with electroencephalography (EEG) and event-related potential (ERP) reflections of attentional bias and disengagement.

When an event occurs, large groups of neurons are activated synchronously, resulting in postsynaptic potentials that can be detected by EEG at specific times when time-locked to the event (Kappenman and Luck 2012). Therefore, EEG allows us to distinguish brain activity associated with the onset of attentional bias (cue-related activity) from brain activity related to the outcome of this bias (target-related activity). For attentional bias, the sequence of ERPs consists of the Early Directing Attention Negativity (EDAN) and the Late Directing Attention Positivity (LDAP) indices of parietal activity. It also includes the Anterior Directing Attention Negativity (ADAN) of frontal activity (Van Der Lubbe et al. 2006). The outcome of bias was identified as the modulation of target-induced parietal P1 and N1 peaks by validity. More specifically, P1 and N1 are increased for validly cued targets compared to invalidly cued targets (Mangun and Hillyard 1991; Meinke et al. 2006). Similarly, attentional disengagement is thought to be reflected in the modulation of the Late Positive Deflection (LPD) (Mangun and Hillyard 1991; Meinke et al. 2006) in the areas that drive inhibitory control, including the right inferior Frontal Gyrus (rIFG) and temporal parietal junction (Corbetta et al. 2008), and it is larger to invalidly cued targets than validly cued targets. Previous studies show that stop signals (assessed by the Stop Signal Task (SST)) evoke a positivity of approximately 300 ms Latency (P300) that is modulated by successful inhibition (Lansbergen et al. 2007). Hence, it was argued that the LDP consists of P300 (Mangun and Hillyard 1991): It is also enhanced when a specific stimulus requires attentional disengagement.

An ideal way to investigate these potential indices of self-regulation is to manipulate frontal asymmetry directly and to assess its effects on them (i.e., FAA, ERPs, and RTs). We utilized tDCS, which delivers low-level electrical currents through the scalp to change cortical activity in expected regions of the brain (Schestatsky et al. 2013). Results of a recent study suggest that tDCS over DLPFC affects self-regulation (Kekic et al. 2017).

To the best of our knowledge, prior studies have not taken into account the impact of tDCS on FAA, as a potential index, to explore whether the effects of tDCS can be attributable to alterations in frontal brain activity asymmetry. We hypothesized that the application of active tDCS would result in heightened right frontal brain activity relative to the left hemisphere. As a result, we anticipated lower FAA scores at frontal sites (F4-F3). We further anticipated that the implementation of active tDCS would lead to a reduction in attentional bias in the reward context compared to the neutral context. This reduction would be evident through a decreased validity effect on response time in the VSC Task, as well as reduced LDAP, N1, and P1 modulation by validity. Simultaneously, we expected active tDCS to enhance attentional disengagement, also reflected by the reduced validity effect on response time in the VSC Task, as well as a stronger LPD response to invalid relative to validly cued targets.

## 2. Methods

### 2.1. Participants

The rationale for the sample size was based on a within-subject pilot study (n = 10), where we estimated the effect size of tDCS concerning FAA as the primary measure. Specifically, using G*Power (Faul et al. 2007, 2009) with a defined power of 80 percent, alpha at .05, and an estimated FAA test-retest correlation of 0.6, we determined that a sample size of 30 in the active intervention group would allow us to detect an effect of f > 0.237 or ηp2 > 0.053. Eligible participants were required to be at least 18 years old and had no psychological or psychiatric disorders, frequent headaches or migraines, metallic implants, epilepsy, significant head trauma in the past, recent head trauma, pacemaker, chronic skin conditions, current drug use, or low command of English. During the main study, some participants withdrew due to various reasons, such as boredom and fatigue. Consequently, a total of 65 adults (46 females) aged 18 to 58 (M_(age)_ = 23.93; SD_(age)_ = 6.08) were recruited, 33 of whom were in the active tDCS group, through social media ads and university courses. Among them, fifty-seven were right-handed. All of the participants had at least a B1 level in English, based on Common European Framework of Reference for Languages - Self-assessment Grid (*Common European Framework of Reference for Languages: Learning, Teaching, Assessment* 2001). These standard demographic questions were implemented in Psytoolkit (Stoet 2010, 2017). Before participating, all individuals provided written informed consent. Each participant received either a voucher or course credit for participation. The study received approval from the Research Ethics Committee at Eötvös Loránd University and was conducted following the principles of the Declaration of Helsinki and its later amendments.

### 2.2. Visuospatial cueing task (VSC)

The VSC task was presented using OpenSesame (Mathôt et al. 2012). It was an adapted version from the original (Posner et al. 1980) and included an intrinsic reward condition based on previous research (Houben et al. 2014; Tsegaye et al. 2022) in addition to the neutral condition. Figure 1A illustrates the details of the task. Trials were randomized, and the condition order (neutral/reward) was counterbalanced across the participants. Only the first keyboard response to a target was logged for each trial. There were four main blocks (two for each condition) with a short practice block at the beginning of the task, and it took approximately 45 minutes in total. Participants were seated approximately 65 cm from the screen. The task began with clear instructions and verbal explanations when needed. Throughout the task, participants were instructed to keep their eyes fixed on the center of the screen. First, a fixation dot was presented for 2000 ms. Subsequently, a central cue (width: 100 (2.5°), height: 100 (2.5°)) was presented for 400 ms. The cue indicated to which side of the screen the participant had to direct their attention. In the case of neutral cues, the cues did not indicate either the right or left side of the screen. Following that, a target was presented at either the left or right visual hemifield, which could be non-cued (presented after the neutral cue), valid (congruent with the location as indicated by the cue), or invalid (incongruent with the location indicated by the cue). The target was bar-shaped and could be either short (width: 100 (2.56°), height: 120 (3.07°)) or long (width: 100 (2.56°), height: 200 (5.12°)). The required response depended on the height of the bar, and stimulus-response assignments were counterbalanced across the participants. The intertrial interval was set to 3100 ms. The neutral and reward contexts differed in terms of the targets. Targets in the neutral condition consisted of solid grey bar-shaped targets. In the intrinsic reward condition, the bar-shaped targets represented palatable food images, such as chips, chocolate, cookies, and cashew nuts. These images are easy to recognize and are high in fat and sugar. Attentional bias was computed by subtracting the mean RT to valid targets from non-cued targets. A larger value indicates a stronger attentional bias. Attentional disengagement was computed by subtracting the RT to non-cued targets from invalidly cued targets. A larger value indicates lower attentional disengagement.

**Figure 1.**
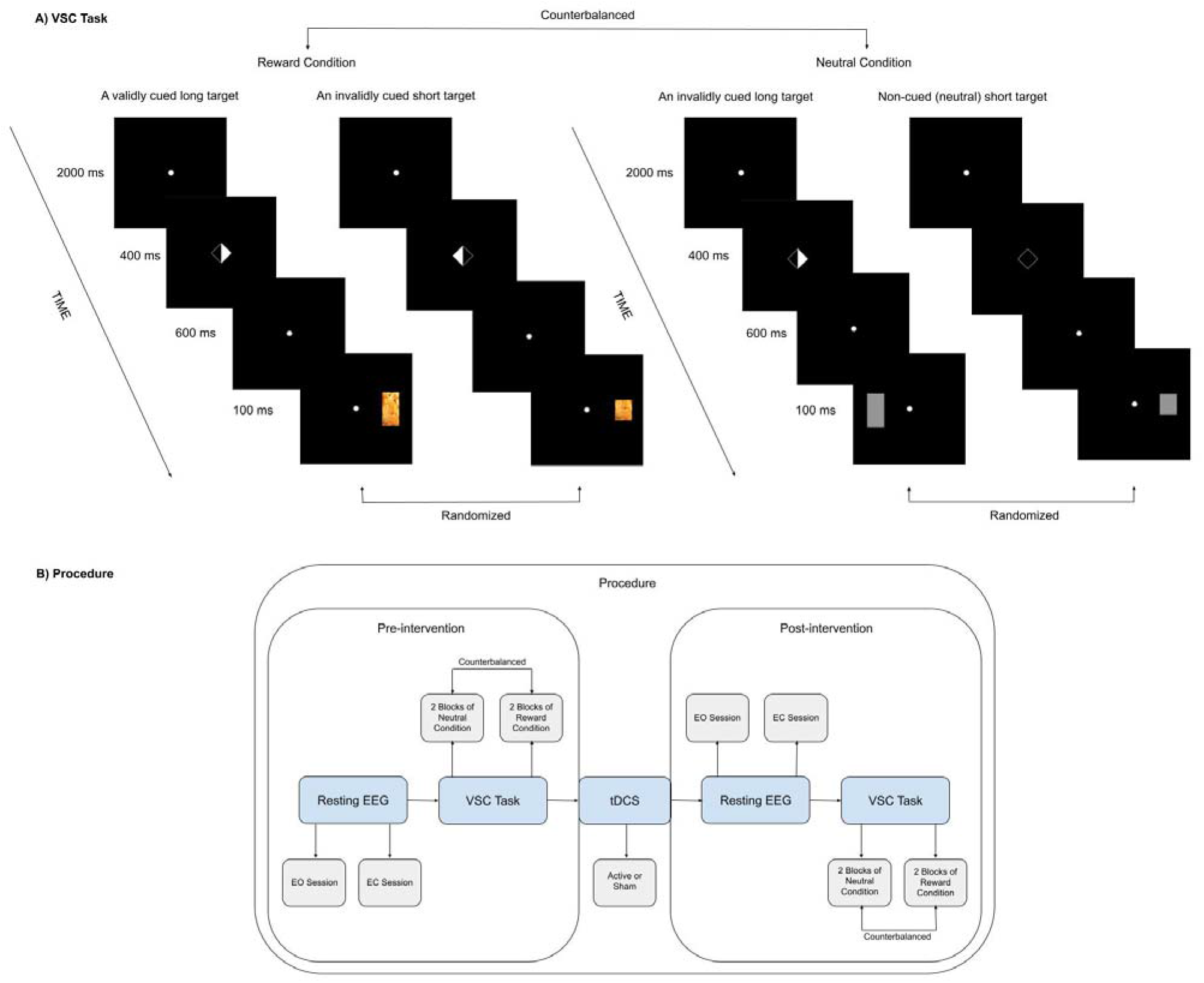
A) This figure illustrates four example trials in the VSC task. Depending on the counterbalanced order, participants began the VSC Task in either the neutral or reward condition. The first two trials (on the left) display a validly cued long target and an invalidly cued short in the reward condition, respectively. The last two trials (on the right) represent the neutral condition. The last trial depicts a non-cued trial in which there is no cue indicating the potential location of a target, and a short neutral target is displayed on the right hemifield. The target sizes overall, and the target types in the reward condition (i.e., chocolate, cookies, nuts, and cashews) were randomized. B) The procedure begins with collecting pre-intervention resting state EEG data. It was collected in two sessions, one with 5 minutes of eyes open (EO) and another with 5 minutes of eyes closed (EC) conditions. Subsequently, participants performed the VSC task. Each participant received both the neutral and reward conditions in a counterbalanced order. Following that, either active or sham tDCS was administered. For further details regarding the intervention, please see Figure 2. After the tDCS intervention, we repeated the same procedure for the post-modulation assessment, which included the resting-state EEG and the VSC Task.

### 2.3. EEG recording

Scalp voltages were recorded using the NeXus-32 from Mind Media (“Nexus-32” n.d.), employing a 21-channel cap Ag/AgCl electrodes placed according to the 10-20 system. The sampling frequency was 512 Hz, and activity was referenced to linked-ears. The horizontal electrooculography (HEOG) was recorded bipolarly from the outer canthi of both eyes, and the vertical EOG (VEOG) was recorded supra-relative to infra-orbital. EEG was recorded continuously except during tDCS-application.

### 2.4. Frontal alpha asymmetry (FAA)

FAA scores were calculated based on resting EEG states collected during 5 minutes of eyes open (EO) and 5 minutes of eyes closed (EC) sessions before and after the intervention. By comparing EEG resting-state recordings in these two conditions, it is easier to understand the impact of sensory input, internal cognitive processes, and the brain’s intrinsic activity, leading to a comprehensive understanding of the brain’s functional organization in different states (Barry et al. 2007). The EEG data was processed using BrainVision Analyzer 2 (www.brainproducts.com). Congruent with previous reports (Smith et al. 2017), FAA was calculated as follows. First, we applied a high-pass filter of 0.5 Hz, a low-pass filter of 40 Hz, and a 50 Hz notch filter. Then, we excluded the initial and final 10 seconds of each approximate 5-minute segment to account for potential artifacts. These segments were further divided into 2-second epochs. The epochs of the EO condition were corrected for ocular artifacts based on VEOG and HEOG electrodes using independent component analysis (ICA). The segments were baseline corrected using the −100 – 0 ms interval and whole-segment baseline corrected. Epochs containing any remaining artifacts (based on 75 µV max amplitude +/− relative to baseline criterion) were discarded, and Power Spectral Density (PSD) was calculated using Fast Fourier Transform FFT with a 10% Hanning window. Afterwards, the epochs were averaged and mean activity in the alpha frequency band (8-13 Hz) was calculated, and values were exported for the relevant electrodes. Alpha power was corrected for skew via natural log transform (Smith et al. 2017) using SPSS 22 (IBM Corporation n.d.). Lastly, frontal asymmetry was calculated by subtracting the mean of log-transformed alpha at lateral left electrode sites from right electrode sites (F4-F3 and F8-F7). Thus, a higher FAA value reflects higher left frontal activity (i.e., lower left frontal alpha power).

### 2.5. Event-related potentials (ERPs)

First, following previous studies (Logemann et al. 2014a, 2014b), EEG data were filtered (offline) using a low pass filter of 30 Hz, a high pass filter of 0.1592, and a notch filter of 50 Hz. Data were segmented into epochs ranging from −100 ms to 2300 ms cue-locked and from −100 to 1000 ms target-locked. Epochs were corrected for blinks (VEOG) using the Gratton et al., (Gratton et al. 1983) algorithm. Segments with evidence of horizontal eye movement higher than 60 µV were rejected (Van Der Lubbe et al. 2006). Subsequently, epochs were baseline-corrected with the baseline set at −100 to 0 ms. The cue-locked ERPs indicated previously were calculated in accordance with van der Lubbe et al. (Van Der Lubbe et al. 2006). The activity at ipsilateral electrodes was subtracted from the activity at contralateral electrodes relative to the cue direction for each participant. Based on van der Lubbe et al. (Van Der Lubbe et al. 2006), the EDAN and LDAP were quantified in the 240-280 ms and 560-640 ms time window at electrode pairs P3/P4 and T3/T4, respectively, while the ADAN was quantified in the 400-480 ms at electrode pair F7/F8. However, after visual inspection of grand average waveforms, the latency of the 240-280 ms time window for the EDAN was adjusted to 190-250 ms while the ADAN time window 400-480 ms was adjusted to 320-400 ms. Based on Mangun and Hillyard (Mangun and Hillyard 1991), the P1 and N1 were analyzed at parietal electrodes P3 and P4 and quantified as the mean amplitude in time windows 100-137 ms and 141-188 ms, respectively, while the LPD was quantified as the mean amplitude in 229-299 ms time window at Cz electrode (Mangun and Hillyard 1991). The P1, N1, and LPD *effects* were computed by subtracting the mean activity within the aforementioned time windows relative to invalidly cued target onset from the activity relative to validly cued target onset.

### 2.6. Transcranial direct current stimulation (tDCS)

Brain modulation aimed to increase the activity of the right relative to the left dorsolateral prefrontal cortex. Direct electrical current was transmitted using a pair of saline-soaked, circular sponge electrodes (25 cm^2^) and delivered via STARSTIM-8, Neuroelectrics (www.neuroelectrics.com). The anode was placed on the right DLPFC (F4), while the cathode was placed on the left DLPC (F3), based on the 10–20 system. A constant current of 2 mA was applied for 20 minutes (Kelley et al. 2017) (Fig.2) unless it was a sham condition. In the sham condition, the same electrical current was only briefly applied at the beginning of the session to generate a similar experience. The employed tDCS procedures have been demonstrated to be safe in healthy participants (Iyer et al. 2005).

**Figure 2.**
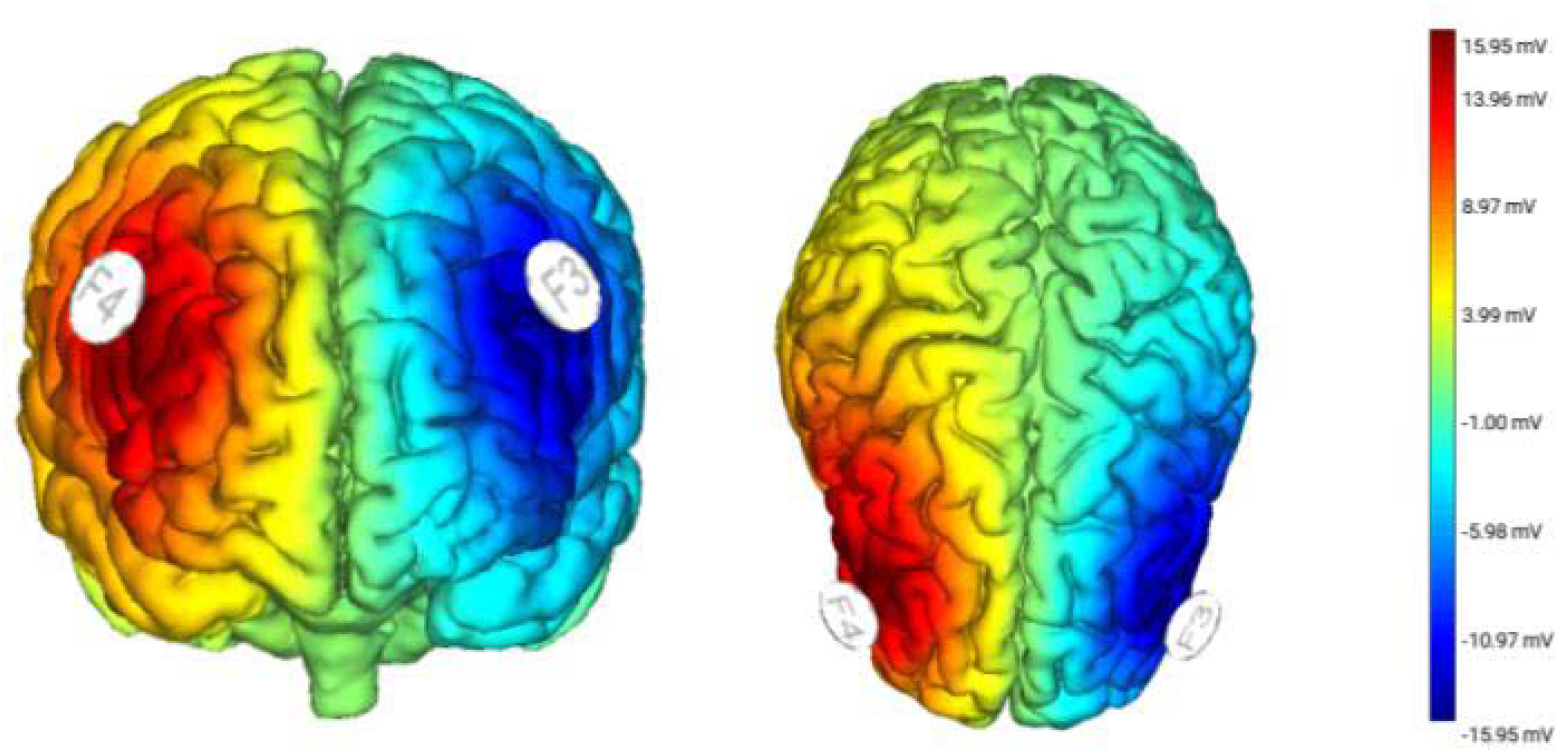
This figure shows the placement of the tDCS electrodes and measured electric fields from the front and top, respectively. A constant current of 2 mA was applied via two circular sponge electrodes (25 cm^2^), placed over positions F4 (anode) and F3 (cathode) with saline solution. The maximal electric field strength was 15.95 µV for the anodal electrode.

### 2.7. Procedure

We employed a randomized, triple-blind, sham-controlled design. We considered within-subject factors, including time (pre-/post-intervention) and condition (neutral/reward), as well as a between-subject factor, which was the group (active/sham tDCS). We preregistered our detailed research plan based on our pilot study on Open Science Framework (https://osf.io/5hvpw?view_only=b528cd416ff047e8be789bc4da284f36). When a participant arrived at our laboratory, they read the information letter, checked the exclusion criteria, and signed the informed consent form. Subsequently, they were seated in a comfortable chair in a dimly lit testing room for the placement of EEG electrodes on the scalp. After placing the electrodes, we collected resting EEG data, with blocks of both EO and EC sessions. Following the resting EEG recording, participants completed the questions. Subsequently, they proceeded to the first part of the VSC Task before the tDCS intervention. EEG was recorded throughout the resting states and computer tasks, but not during the stimulation as the cap was changed. Subsequently, they were randomly assigned to receive either active or sham tDCS. The tDCS software provided by Neuroelectrics (www.neuroelectrics.com) completed the assignments, ensuring that the individuals involved, including the participants, experimenters, and data analysts, remained unaware of who was actually receiving the stimulation. After the tDCS intervention, we conducted the same procedure again for the post-modulation assessment, including the resting-state EEG and the VSC Task. Please consider Figure 1B for more details. The pre-/post-assessments and the intervention all took place on the same day, lasting approximately five hours.

### 2.8. Statistical analyses

The statistical analyses were conducted using SPSS 22 (IBM Corporation n.d.) and R (“R software: a tool analysing experimental data” 2016). First, participants with missing values and outliers exceeding 3 standard deviations (SD) from the mean were excluded in case of evident erroneous data. Then, we used repeated measures ANOVA (2×2×2) to test the hypotheses. The electrophysiological variables under investigation included FAA (EO/EC), ADAN, EDAN, LDAP, N1 *effect,* P1 *effect*, and LPD *effect*, as well as the behavioral indices of attentional bias and disengagement. For all analyses, the significance level was set at 0.05. We also performed a series of Bayesian repeated measures ANOVA for the null results. They were used with Bayes factor 01 (BF_01_), which is in favor of null hypotheses over alternative hypotheses. More particular, a score of less than 0.33 indicates strong support for alternative hypotheses (Wagenmakers et al. 2011, 2018). Secondary analyses were performed on the behavioral and ERP indices of attentional processes, excluding participants who showed an absent validity effect at baseline in the VSC task. Additionally, exploratory analyses were conducted to explore the effects of time and group in the reward condition. Results from exploratory and Bayesian analysis are included in the supplementary materials.

## 3. Results

### 3.1. Did the tDCS have an impact on frontal alpha asymmetry?

To test whether the tDCS intervention affected the primary target resting-state FAA, a series of repeated measures of ANOVA with the within-subjects factor of time (pre-/post-intervention) and the between-subject factor of group (active/sham tDCS) was conducted. The frequentist and Bayesian analyses showed that there is no effect of time and group interactions on FAA F4-F3 (EO): F(1, 114) = 0.28, p = 0.594, η_p_^2^ = 0.002, BF_01_ > 0.3 and on FAA F4-F3 (EC): F(1, 114) = 0.49, p = 0.484, η_p_^2^ = 0.004, BF_01_ > 0.3. These results were previously reported by another study in detail (Akil et al. 2023).

### 3.2. Did the tDCS affect the behavioral indices of visuospatial attention?

We conducted a series of repeated measures ANOVA with the within-subject factors of time (pre-/post-intervention) and condition (neutral/reward), as well as the between-subject factor of group (active/sham tDCS). The statistical results are provided in Table 1. Although there was a significant main effect of condition on attentional disengagement, F(1, 232) = 8.59, p = 0.003, η_p_^2^ = 0.035, BF_01_ < 0.3, its interaction with time and group factors was not significant F(1, 232) = 0.01, p = 0.909, η_p_^2^ = <0.001, BF_01_ > 0.3. On the other hand, the condition + group model was also in favor of the alternative hypothesis, BF_01_ < 0.3. Please refer to Tables 6 and 7 in the supplementary materials for further information. The direction of the relationship between attentional disengagement and condition is illustrated in Figure 3. The figure revealed reduced attentional disengagement in the reward condition compared to the neutral condition.

**Figure 3.**
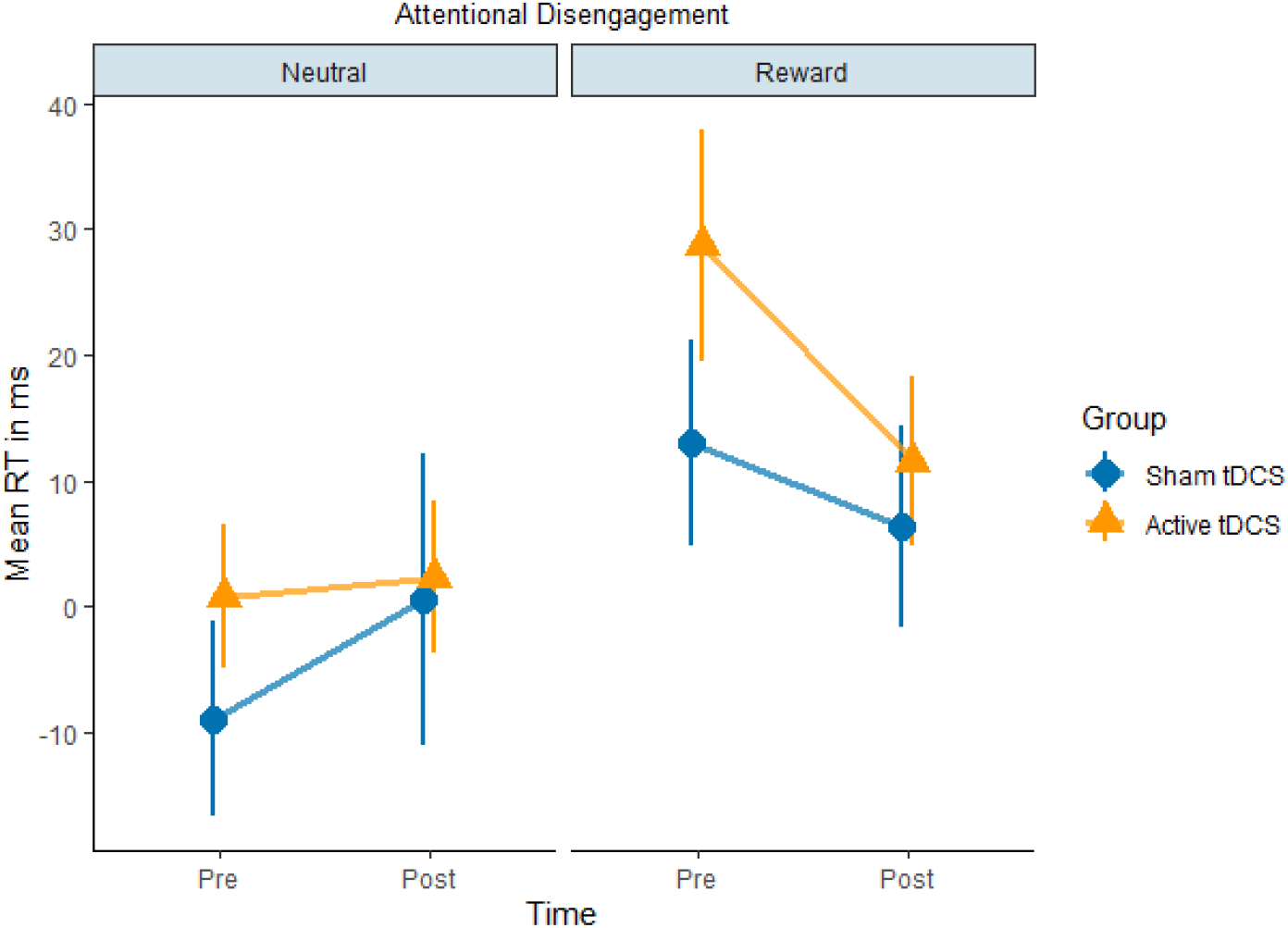
This figure shows the mean RTs of attentional disengagement based on time (pre-/post-intervention), condition (neutral/reward), and group (sham/active tDCS) factors. It depicts reduced attentional disengagement in the reward condition compared to the neutral condition. Please note that attentional disengagement was calculated by subtracting the mean RT to non-cued targets from invalidly cued targets. Therefore, a greater value indicates lower attentional disengagement.

**Table 1.** Results of attentional bias and attentional disengagement models.

| Variables | Df | F | P | $\eta_p^2$ |
| --- | --- | --- | --- | --- |
| **Attentional Bias (N = 60)** |  |  |  |  |
| Time | 1 | 0.01 | 0.917 | <0.001 |
| Condition | 1 | 2.60 | 0.108 | 0.011 |
| Group | 1 | 2.05 | 0.153 | 0.008 |
| Time x Condition | 1 | 0.18 | 0.670 | <0.001 |
| Time x Group | 1 | 1.22 | 0.270 | 0.005 |
| Condition x Group | 1 | 0.03 | 0.851 | <0.001 |
| Time x Condition x Group | 1 | 0.11 | 0.730 | <0.001 |
| Error | 232 |  |  |  |
| **Attentional Disengagement (N = 60)** |  |  |  |  |
| Time | 1 | 0.42 | 0.517 | 0.001 |
| Condition | 1 | 8.59 | 0.003* | 0.035 |
| Group | 1 | 2.05 | 0.153 | 0.008 |
| Time x Condition | 1 | 2.44 | 0.119 | 0.010 |
| Time x Group | 1 | 0.67 | 0.413 | 0.002 |
| Condition x Group | 1 | 0.17 | 0.677 | <0.001 |
| Time x Condition x Group | 1 | 0.01 | 0.909 | <0.001 |
| Error | 232 |  |  |  |

Note: Participants with missing values and erroneous data were excluded from the analyses.

### 3.3. Did the tDCS affect cue-locked event-related potentials?

A series of repeated measures ANOVA was conducted on each cue-locked ERP. The within-subject factors included time (pre-/post-intervention) and condition (neutral/reward), while the between-subject factor was group (active/sham tDCS). The results concerning the EDAN, ADAN, and LDAP can be observed in Figures 4, 5, and 6, respectively. For detailed inferential statistics, refer to Table 2. We found that the effect of time, condition, and group interactions on LDAP showed a trend towards significance, F(1, 56) = 2.98, p = 0.090, η_p_^2^ = 0.051, BF_01_ > 0.3. There was no effect of time, condition, and group interactions on the EDAN and ADAN. These results were supported by Bayesian analyses and they can be found in Tables 8, 9, and 10 in the supplementary materials.

**Figure 4.**
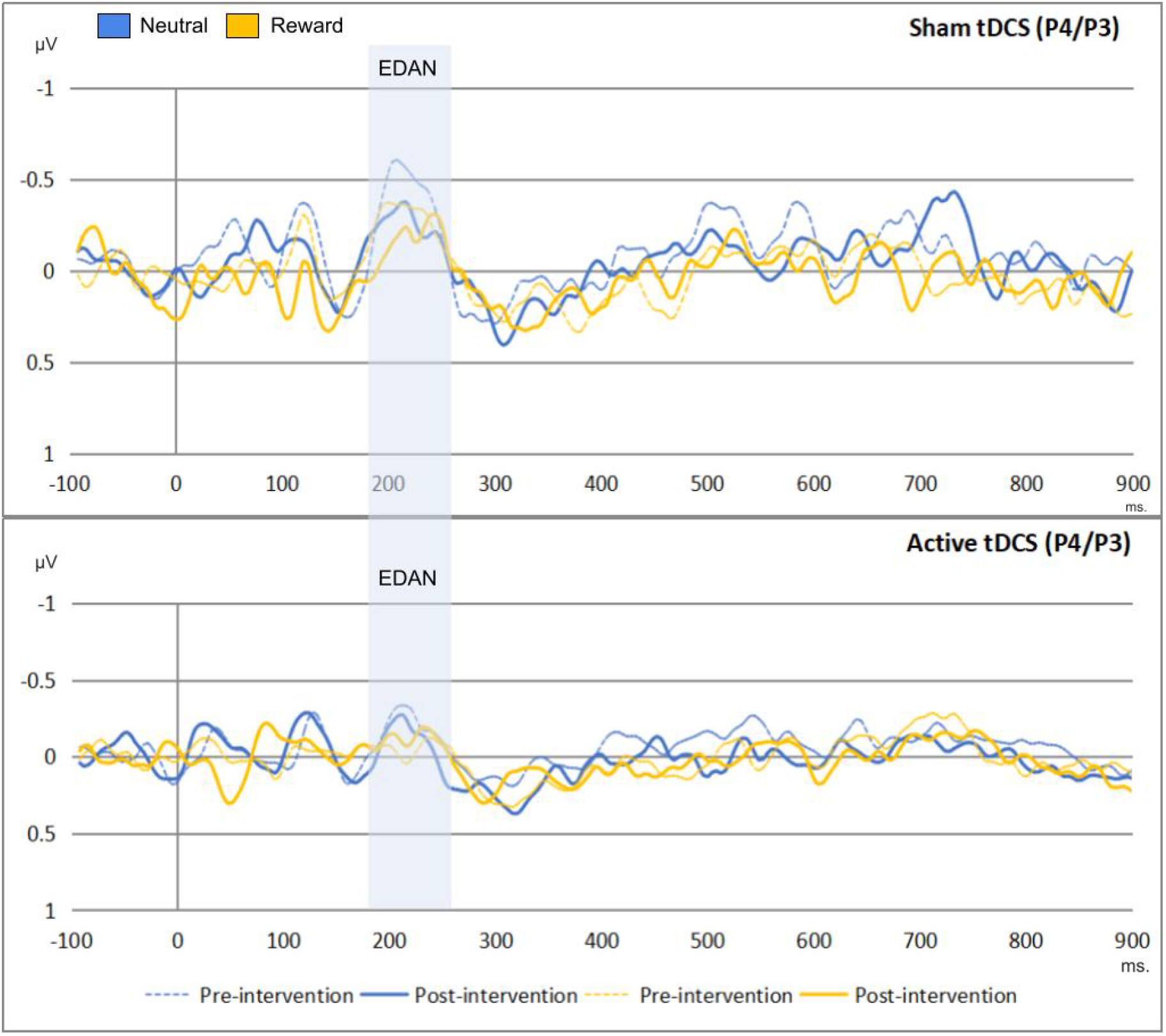
This figure shows the effects of time, condition, and group on the EDAN (190-250 ms). The x-axis represent the time in milliseconds, while y-axis represents the EDAN scores in microvolts. Based on the depiction, the EDAN was replicated in the study regardless of time, condition, and group effects.

**Figure 5.**
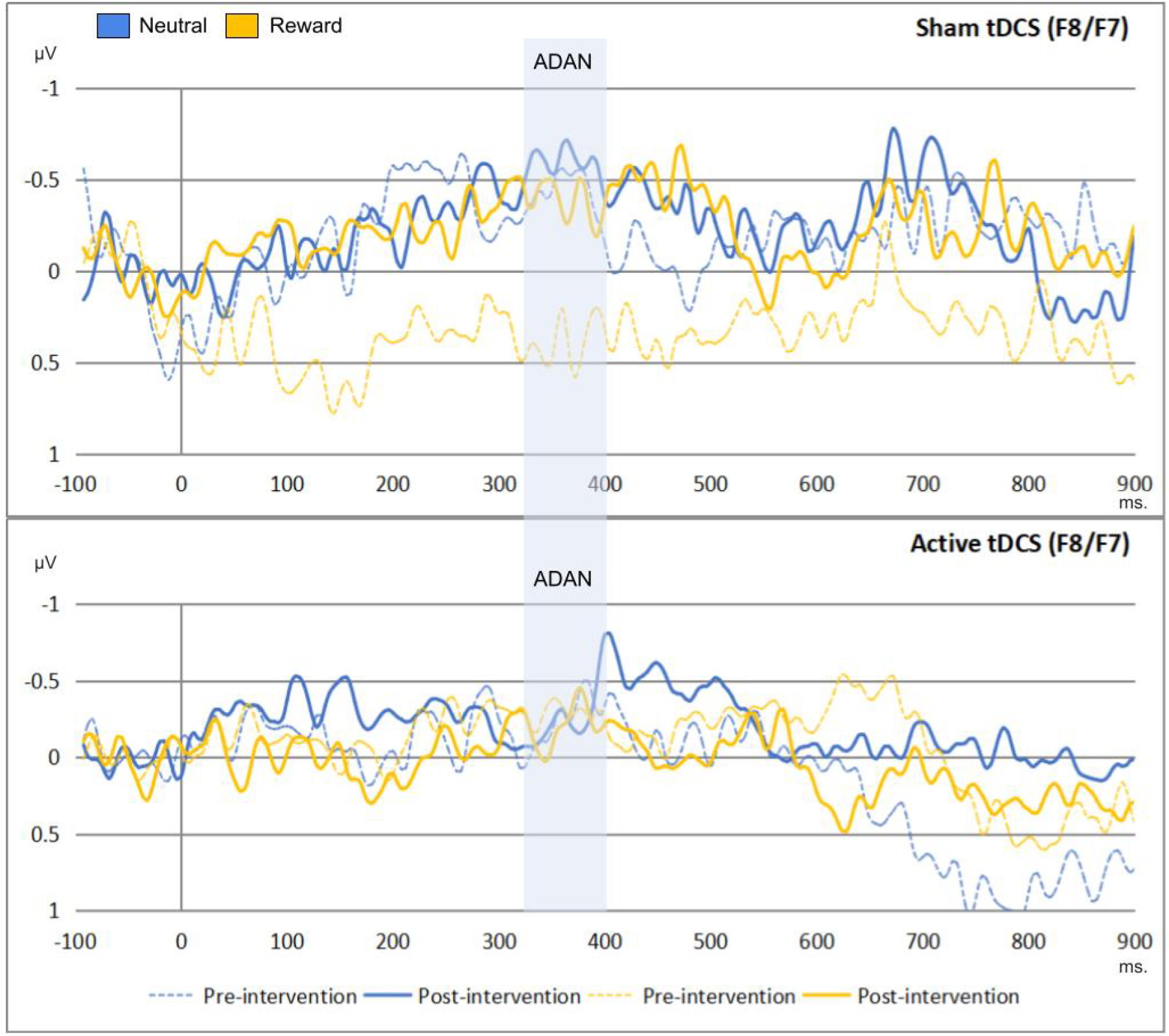
This figure shows the effects of time, condition, and group on the ADAN (190-250 ms). The x-axis represent the time in milliseconds, while y-axis represents the EDAN scores in microvolts. Based on the illustration, the ADAN was replicated in the study even though there were no significant effects of time, condition, and group factors.

**Figure 6.**
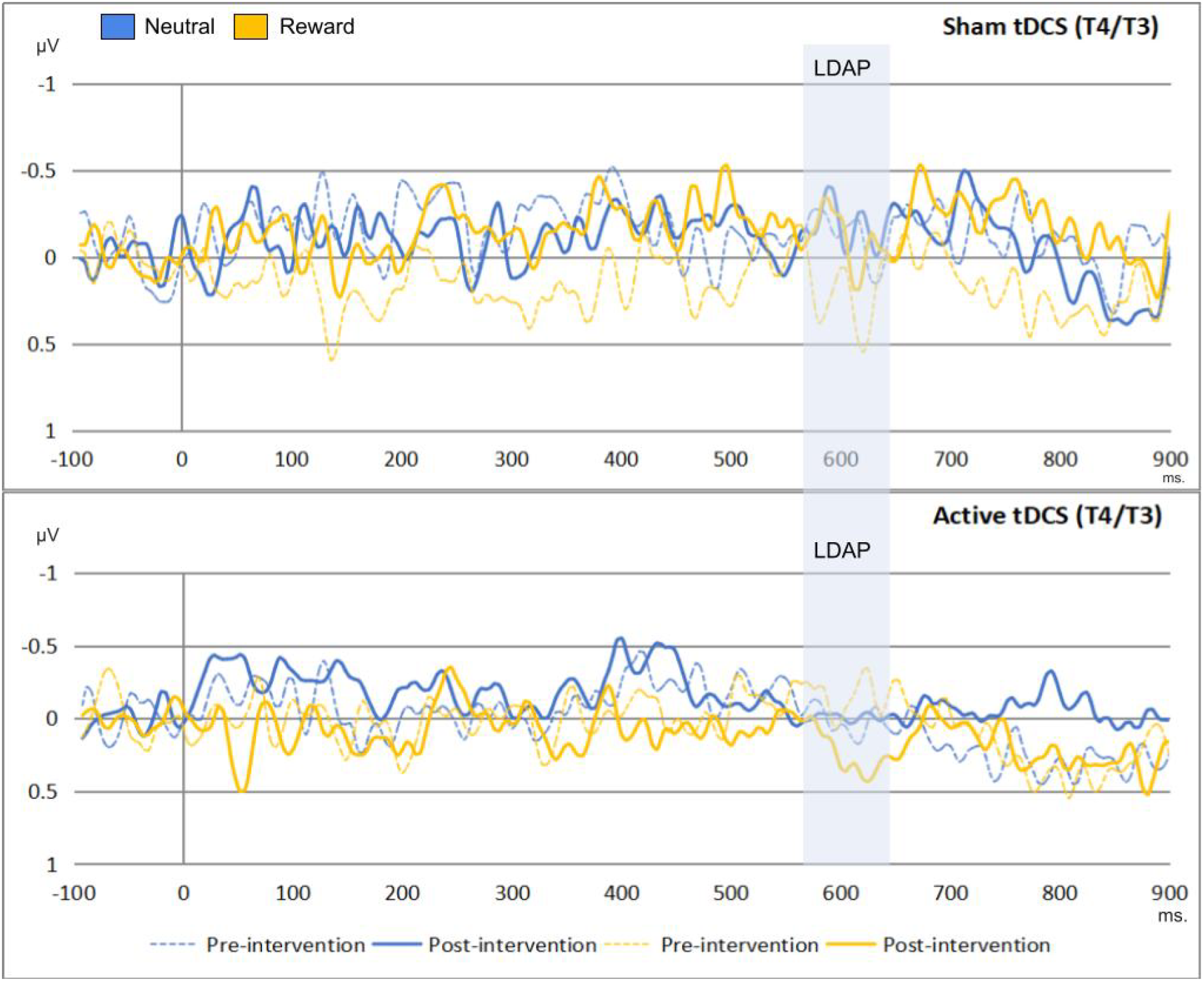
This figure shows the effects of time, condition, and group on the LDAP (560-640 ms). The x-axis represent the time in milliseconds, while y-axis represents the EDAN scores in microvolts. Based on the depiction, the LDAP was replicated in the study.

**Table 2.** Results of cue-locked ERP models.

| Variables | Df | F | p | $\eta_p^2$ |
| --- | --- | --- | --- | --- |
| EDAN at 190-250 ms. (P4/P3) (N = 58) |  |  |  |  |
| Time | 1 | 2.13 | 0.149 | 0.037 |
| Condition | 1 | 0.71 | 0.402 | 0.013 |
| Group | 1 | 0.71 | 0.401 | 0.013 |
| Time x Condition | 1 | 0.69 | 0.408 | 0.012 |
| Time x Group | 1 | 3.09 | 0.084 | 0.052 |
| Condition x Group | 1 | <0.01 | 0.991 | <0.001 |
| Time x Condition x Group | 1 | 0.28 | 0.597 | 0.005 |
| Error | 56 |  |  |  |
| ADAN at 320-400 ms. (F8/F7) (N = 58) |  |  |  |  |
| Time | 1 | 0.92 | 0.762 | 0.002 |
| Condition | 1 | 0.11 | 0.738 | 0.002 |
| Group | 1 | 0.07 | 0.784 | 0.001 |
| Time x Condition | 1 | 1.57 | 0.215 | 0.027 |
| Time x Group | 1 | 0.20 | 0.656 | 0.004 |
| Condition x Group | 1 | 0.86 | 0.356 | 0.015 |
| Time x Condition x Group | 1 | 2.23 | 0.141 | 0.038 |
| Error | 56 |  |  |  |
| LDAP at 560-640 ms. (T4/T3) (N = 58) |  |  |  |  |
| Time | 1 | 0.11 | 0.733 | 0.002 |
| Condition | 1 | 0.01 | 0.896 | <0.001 |
| Group | 1 | 0.02 | 0.874 | <0.001 |
| Time x Condition | 1 | <0.01 | 0.958 | <0.001 |
| Time x Group | 1 | 0.01 | 0.899 | <0.001 |
| Condition x Group | 1 | <0.01 | 0.923 | <0.001 |
| Time x Condition x Group | 1 | 2.98 | 0.090 | 0.051 |
| Error | 56 |  |  |  |

### 3.4. Did the tDCS affect target-locked event-related potentials?

We conducted a series of repeated measures ANOVA to assess whether the tDCS intervention influenced the target-locked ERPs. The within-subject factors for the models were time (pre-/post-intervention) and condition (neutral/reward), while the between-subject factor was group (active/sham tDCS). We found no significant effect of time, condition, and group interactions on target-locked ERPs as shown in Tables 3, 4, and 5 for the P1 *effect*, N1 *effect*, and LPD *effect*, respectively. These results are also illustrated in Figure 7 and Figure 8. Similar to previous explorations, the P1 *effect* was not identified(Logemann et al. 2014b). The models were in favor of the null hypotheses: Please refer to Tables 11, 12, 13, 14, and 15 in the supplementary materials for Bayesian results.

**Figure 7.**
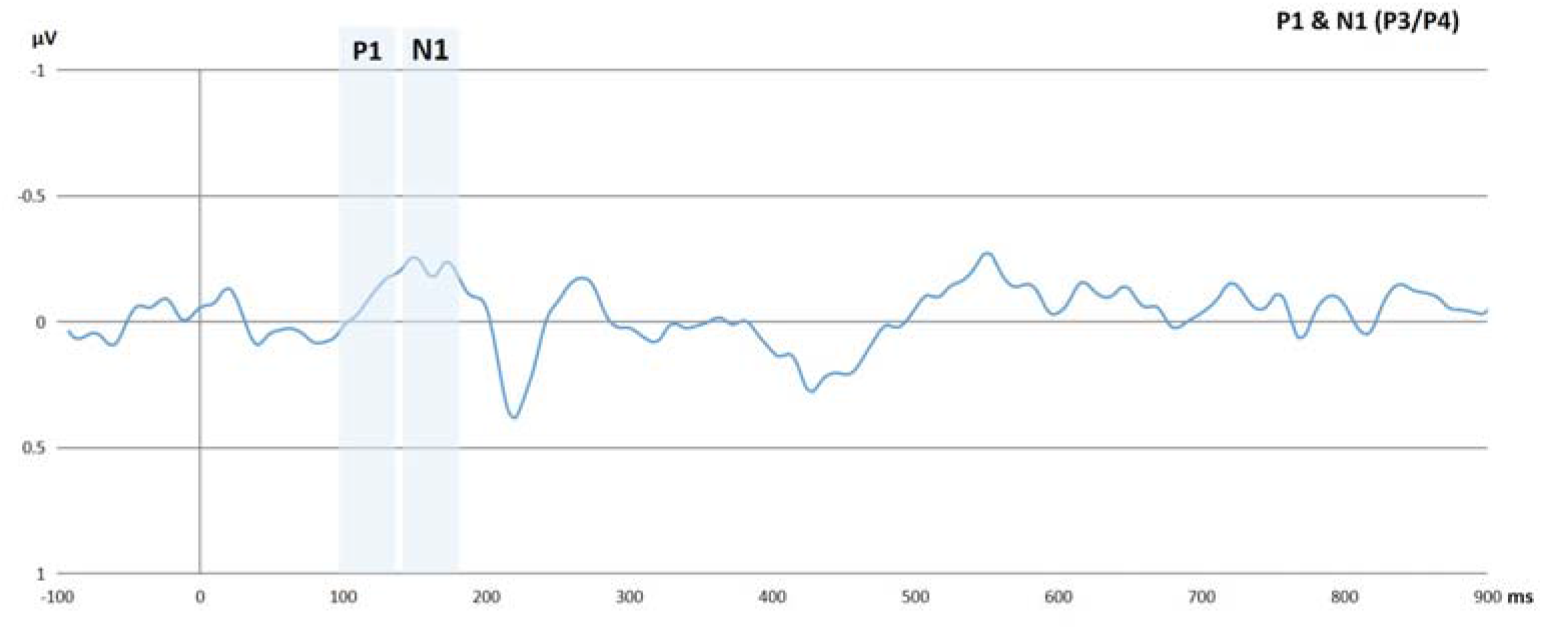
The figure shows the P1 *effect* and N1 *effect*. The x-axis represents the time in milliseconds, while y-axis represents the P1 *effect* and N1 *effect* scores in microvolts. The P1 effect was calculated by subtracting the P1 to invalid trials from the P1 to valid trials. Similarly, the N1 was calculated by subtracting the N1 to invalid trials from the N1 to valid trials.

**Figure 8.**
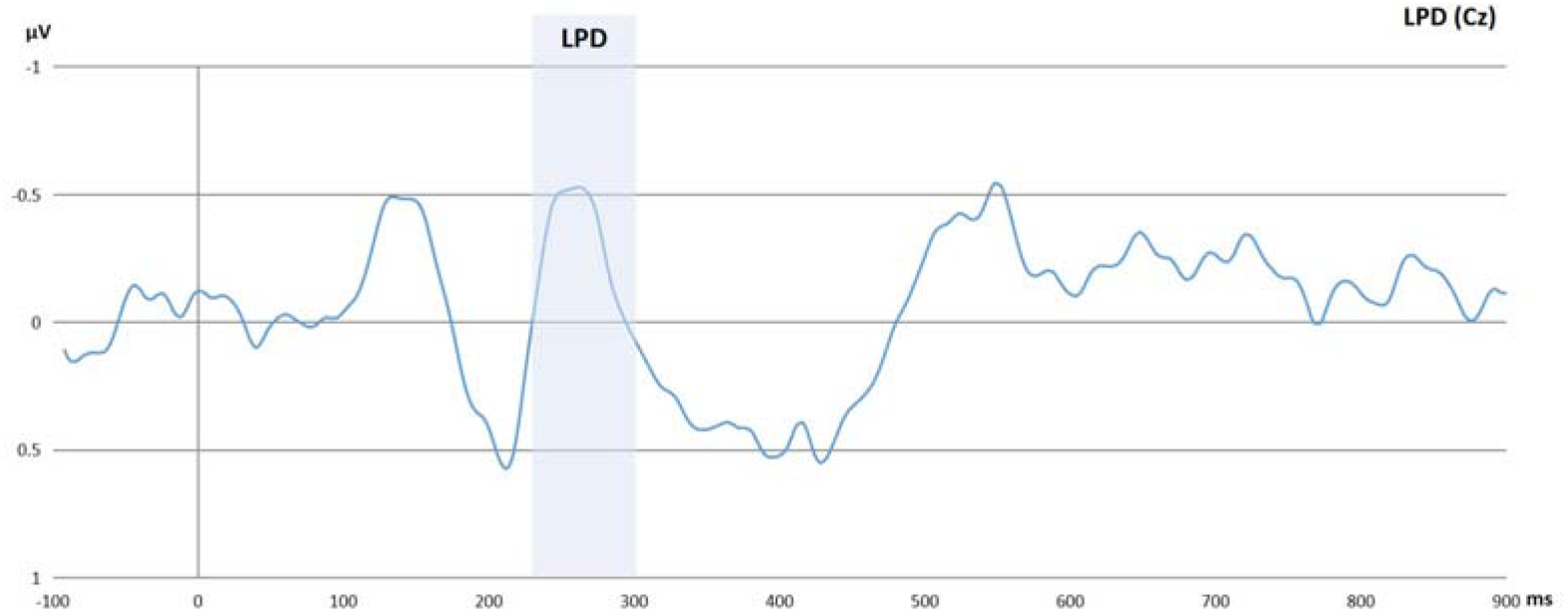
The figure shows the LPD *effect*. The x-axis represents the time in milliseconds, while y-axis represents the LPD *effect* scores in microvolts. The LPD *effect* was calculated by subtracting the LPD to invalid trials from the LPD to valid trials. Therefore, the negative peak represents an enhanced LPD for invalid trials relative to valid trials as well as a reduced LPD for valid trials relative to invalid trials.

**Table 3:** Results of P1 effect models.

| Variables | Df | F | p | $\eta_p^2$ |
| --- | --- | --- | --- | --- |
| **P1 at 100-137 ms. (P3) (N = 55)** |  |  |  |  |
| Intercept |  | 1.25 | 0.269 | 0.023 |
| Time | 1 | 1.85 | 0.179 | 0.034 |
| Condition | 1 | 0.05 | 0.818 | 0.001 |
| Group | 1 | 0.90 | 0.347 | 0.017 |
| Time x Condition | 1 | 1.09 | 0.084 | 0.055 |
| Time x Group | 1 | 0.19 | 0.662 | 0.004 |
| Condition x Group | 1 | 0.82 | 0.368 | 0.015 |
| Time x Condition x Group | 1 | 0.37 | 0.545 | 0.007 |
| Error | 53 |  |  |  |
| **P1 at 100-137 ms. (P4) (N = 55)** |  |  |  |  |
| Intercept |  | 0.79 | 0.375 | 0.015 |
| Time | 1 | 2.60 | 0.113 | 0.047 |
| Condition | 1 | 0.25 | 0.257 | 0.024 |
| Group | 1 | 1.37 | 0.246 | 0.025 |
| Time x Condition | 1 | 2.40 | 0.127 | 0.043 |
| Time x Group | 1 | 1.31 | 0.257 | 0.024 |
| Condition x Group | 1 | 3.88 | 0.054 | 0.068 |
| Time x Condition x Group | 1 | 0.01 | 0.896 | <0.001 |
| Error | 53 |  |  |  |

Note: Participants with missing values and erroneous data were excluded
from the analyses.

**Table 4:**
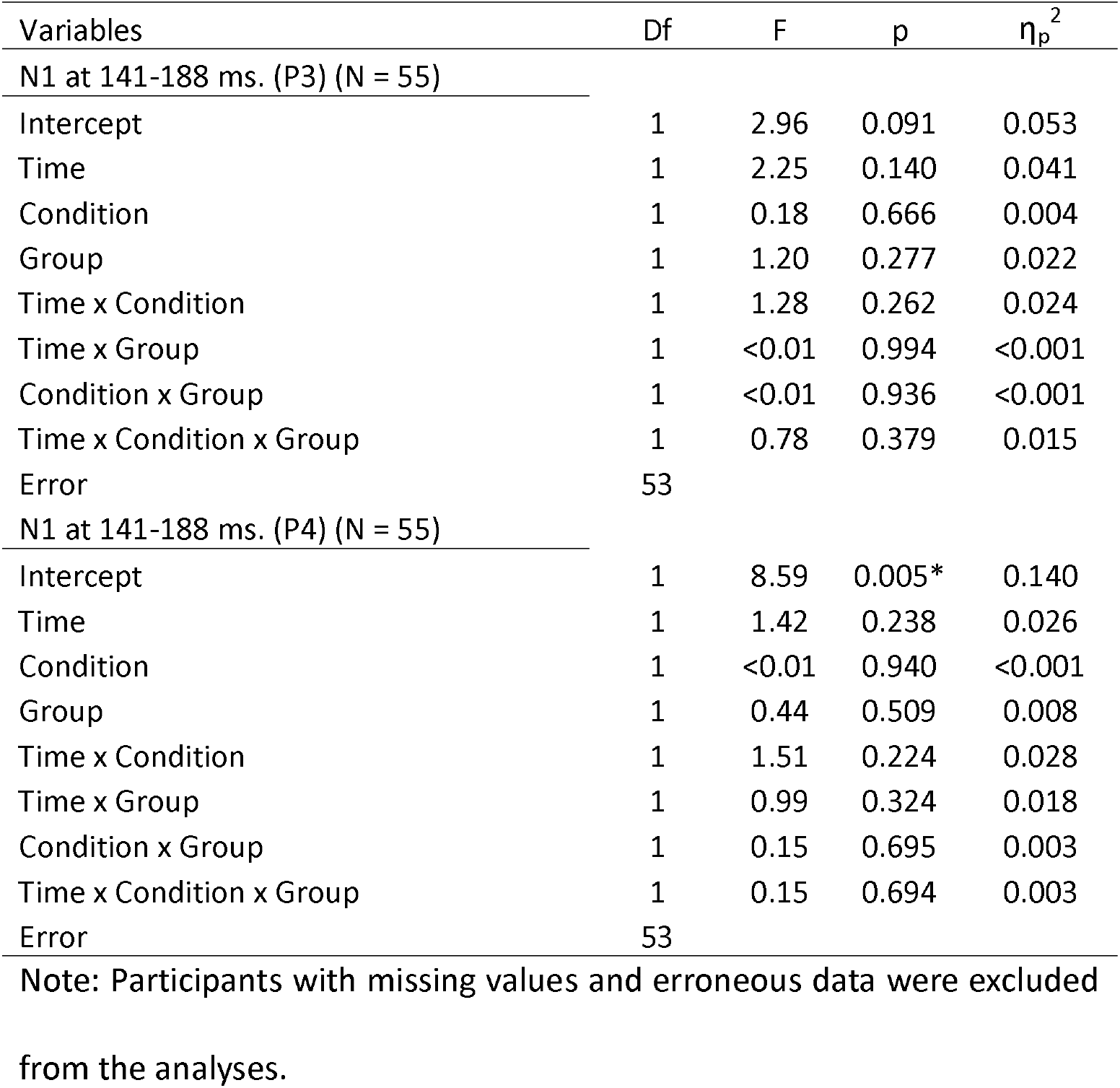
Results of N1 effect models.

| Variables | Df | F | p | $\eta_p^2$ |
| --- | --- | --- | --- | --- |
| **N1 at 141-188 ms. (P3) (N = 55)** |  |  |  |  |
| Intercept | 1 | 2.96 | 0.091 | 0.053 |
| Time | 1 | 2.25 | 0.140 | 0.041 |
| Condition | 1 | 0.18 | 0.666 | 0.004 |
| Group | 1 | 1.20 | 0.277 | 0.022 |
| Time x Condition | 1 | 1.28 | 0.262 | 0.024 |
| Time x Group | 1 | <0.01 | 0.994 | <0.001 |
| Condition x Group | 1 | <0.01 | 0.936 | <0.001 |
| Time x Condition x Group | 1 | 0.78 | 0.379 | 0.015 |
| Error | 53 |  |  |  |
| **N1 at 141-188 ms. (P4) (N = 55)** |  |  |  |  |
| Intercept | 1 | 8.59 | 0.005* | 0.140 |
| Time | 1 | 1.42 | 0.238 | 0.026 |
| Condition | 1 | <0.01 | 0.940 | <0.001 |
| Group | 1 | 0.44 | 0.509 | 0.008 |
| Time x Condition | 1 | 1.51 | 0.224 | 0.028 |
| Time x Group | 1 | 0.99 | 0.324 | 0.018 |
| Condition x Group | 1 | 0.15 | 0.695 | 0.003 |
| Time x Condition x Group | 1 | 0.15 | 0.694 | 0.003 |
| Error | 53 |  |  |  |

Note: Participants with missing values and erroneous data were excluded
from the analyses.

**Table 5:** Results of the LPD effect model.

| Variables | Df | F | p | $\eta_p^2$ |
| --- | --- | --- | --- | --- |
| **LPD at 229-299 ms. (Cz) (N = 55)** |  |  |  |  |
| Intercept | 1 | 6.62 | 0.013* | 0.111 |
| Time | 1 | 3.37 | 0.072 | 0.060 |
| Condition | 1 | 0.34 | 0.559 | 0.006 |
| Group | 1 | 1.26 | 0.267 | 0.023 |
| Time x Condition | 1 | 2.86 | 0.097 | 0.051 |
| Time x Group | 1 | 0.11 | 0.734 | 0.002 |
| Condition x Group | 1 | 1.66 | 0.202 | 0.030 |
| Time x Condition x Group | 1 | 0.14 | 0.703 | 0.003 |
| Error | 53 |  |  |  |

Note: Participants with missing values and erroneous data were excluded
from the analyses.

### 3.5. Exploratory analyses

In line with a previously reported approach (Thiel and Fink 2008), secondary analyses were conducted with individuals who showed a validity effect at baseline in both neutral and reward conditions. As stated previously, the validity effect was calculated based on higher RTs in invalid trials than in valid trials (invalid trials > valid trials). The following analyses were conducted based on this selected sample.

Similar to the previous results, the main effect of condition on attentional disengagement was significant, F(1, 128) = 4.86, p = 0.029, η_p_^2^ = 0.034. Indeed, secondary analyses based on the validity effect did not yield any additional information regarding the behavioral indices of visuospatial attention. Please see Table 16 in the supplementary file for further information.

We also tested the effect of time and group interactions on electrophysiological indices of visuospatial attention in the neutral and reward conditions. Both the LDAP and P1 *effect* were enhanced in the active tDCS group relative to the sham tDCS group in the reward context, F(1, 30) = 6.70, p = 0.015, η_p_^2^ = 0.183 and F(1, 30) = 6.30, p = 0.018, η_p_^2^ = 0.174, respectively. They can also be seen in Figure 9 and 10, respectively. Please also consider Table 19 and 22 in the supplementary file for more information.

**Figure 9.**
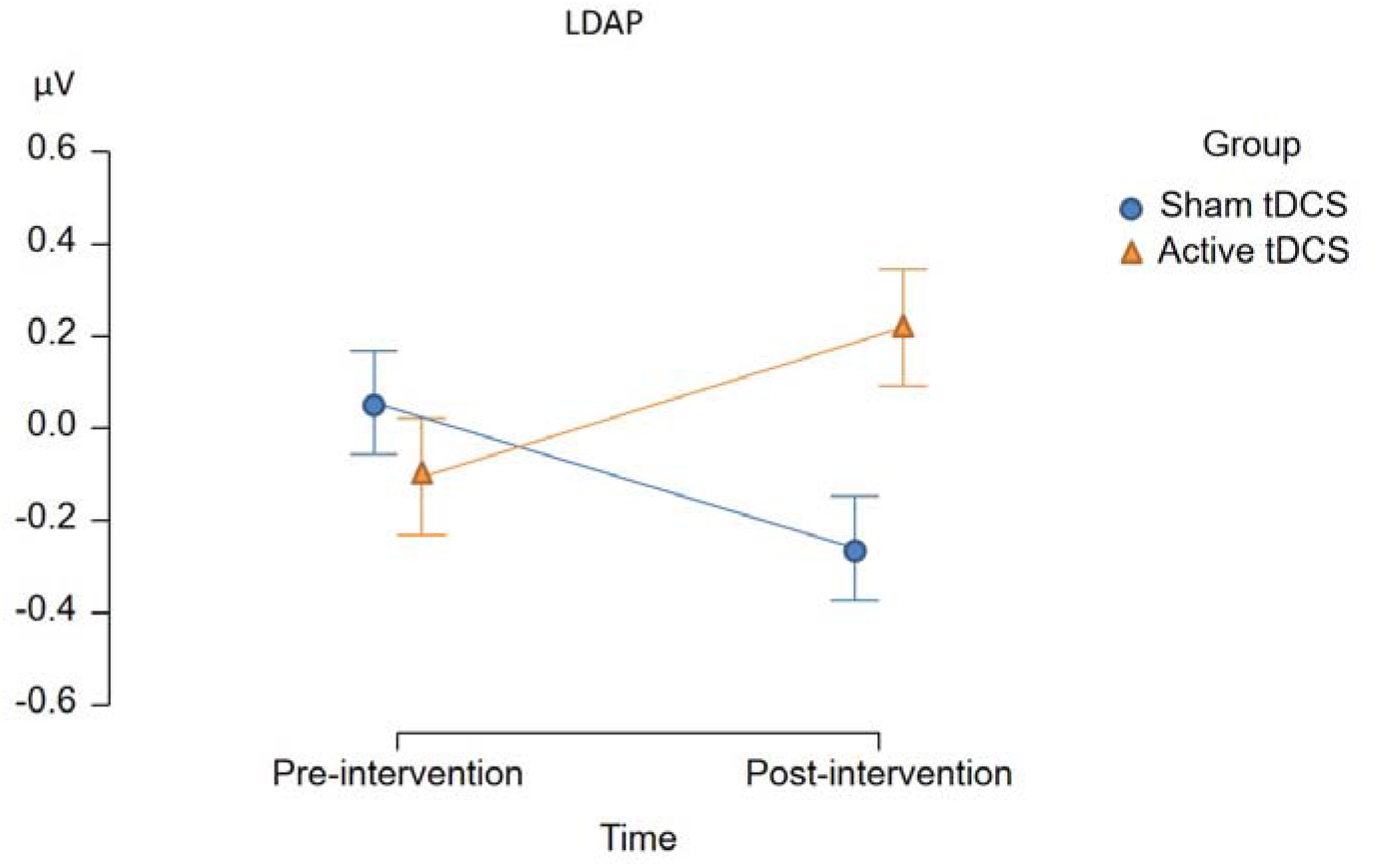
The figure shows the exact effect of time and group interactions on the LDAP in the reward context. The x-axis represents the time factor, while y-axis represents the LDAP in microvolts.

**Figure 10.**
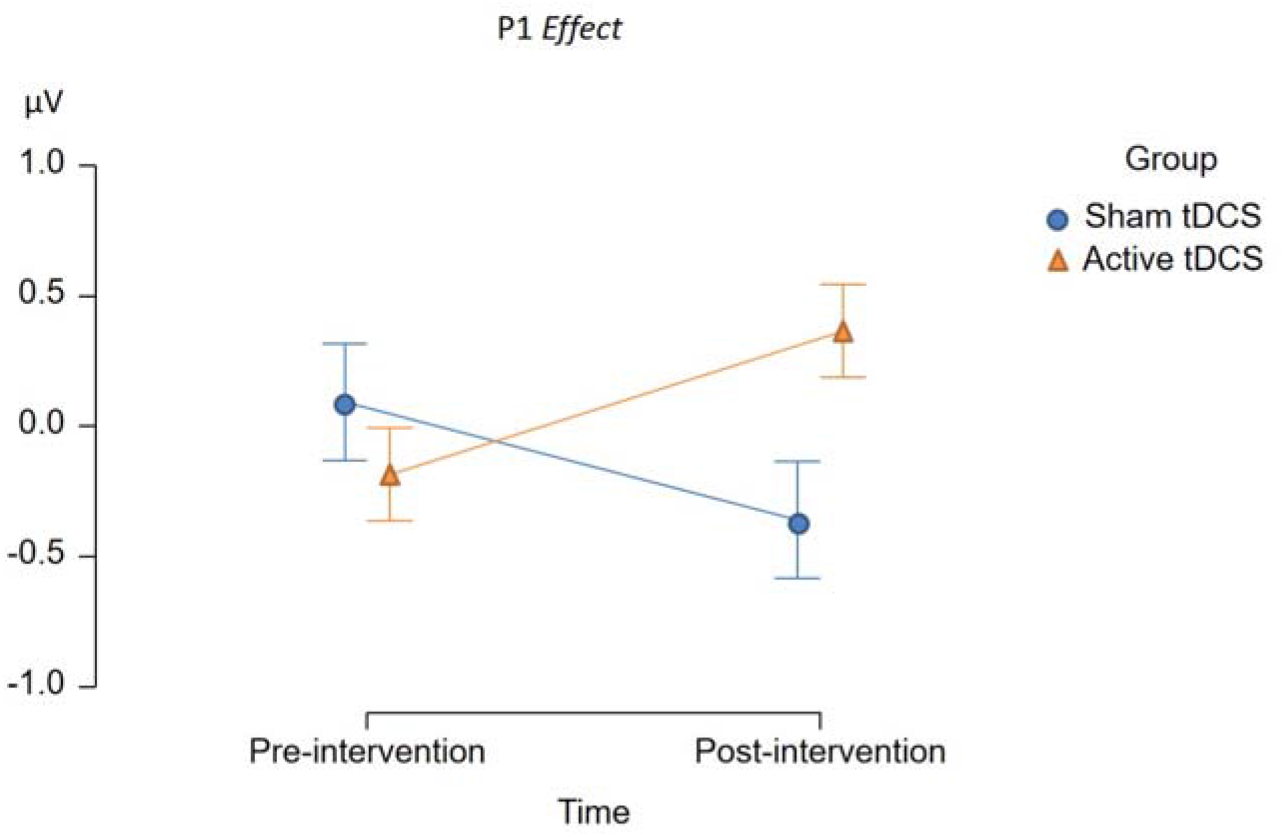
The figure shows the exact effect of time and group interactions on the P1 effect in the reward context. The x-axis represents the time factor, while y-axis represents the P1 effect in microvolts.

## 4. Discussion

The present study investigated the effect of tDCS on FAA as a potential neurophysiological index of self-regulation, as well as on behavioral and brain activity indices of visuospatial attention under the influence of intrinsic reward-related stimuli, specifically palatable food images. The results revealed no significant effect of tDCS on FAA. On the other hand, the exploratory analyses regarding the electrophysiological indices of visuospatial attention demonstrated a statistically significant enhancement of attentional orienting towards food stimuli in the active tDCS group compared to the sham tDCS group. Nevertheless, the effect of tDCS did not translate into observable behavioral changes. We only observed a statistically significant main effect of condition on the RTs, indicating reduced attentional disengagement in the reward condition compared to the neutral condition.

Previous studies suggested that tDCS may affect the asymmetry of frontal brain activity (Kekic et al. 2017). Despite their relevance, these measures are considered indirect. Importantly, to the best of our knowledge, prior studies have not thoroughly investigated whether tDCS actually affects the primary target, which is the asymmetry of frontal brain activity. The results did not show any support for tDCS-induced shifts in the asymmetry of frontal brain activity, as indexed by FAA. Though contrary to the hypothesis, the lack of a significant effect may not be too surprising. The path of electric currents is complex, involving the electrodes, scalp, skull, meninges, and cerebrospinal fluid before reaching the brain. However, each brain has a unique anatomical structure, resulting in different magnitudes and directions of the electric field in each subject (Truong et al. 2013; Laakso et al. 2015; Opitz et al. 2015). Indeed, not only neuronal structure, such as skull thickness, sulcal depth, and gyral depth, can influence the electric field strength, but also there may be “hotspots” that resist electrode positioning (Opitz et al. 2015). Especially when compared to motor cortical tDCS, frontal tDCS produces more variable electric fields (Laakso et al. 2016). This variability may lead to conflicting results in brain stimulation studies (Tremblay et al. 2014). Aside from the variations in individual brain anatomy, the effects of stimulation can also be influenced by factors such as electrode properties, including size and shape, and conductivities, such as gels and saline solutions (Saturnino et al. 2015). Therefore, these findings in the literature suggest that the lack of a significant effect of tDCS on FAA may be attributed to the variability in individual brain structures, the influence of electrode positioning, and the diverse effects of tDCS montages and electrode properties on stimulation outcomes. Future studies should take into account and control for individual differences to ensure the validity and reliability of the findings.

Even though primary analyses did not reveal a significant effect of tDCS on attentional bias and disengagement at the behavioral level, the results indicated a potential effect of tDCS on the electrophysiological measure of attentional orienting, which was confirmed in our exploratory analyses. For these analyses, we employed the same approach as in Thiel et al. (Thiel and Fink 2008) and excluded participants who did not show evidence of cue-induced manipulation of attention. In the resulting sample, tDCS resulted in an enhanced cue-induced LDAP, specifically in the reward context. This can be interpreted as a tDCS-induced enhancement of top-down attentional orienting to spatial locations associated with upcoming reward-associated stimuli. Surprisingly, this effect contradicts the hypothesized shift toward enhanced right versus left frontal activity in our study and previous studies. More specifically, previous research has reported that tDCS enhances inhibitory control (Kekic et al. 2017). The effect of tDCS on enhancing inhibitory control might arguably outweigh its impact on attentional bias and any potential associated response complicating inhibitory control. Therefore, the question arises as to which alternative mechanism fits the observed effect. Likewise, a study found positive effects on inhibitory control irrespective of the stimulation site, pointing towards a more general mechanism: Both anode right/cathode left and anode left/cathode right tDCS increased inhibitory control (Kekic et al. 2017). The answer to these questions can potentially be found in animal studies. Animal studies suggest that tDCS generates a subthreshold depolarization of neuronal membrane potential, leading to an increased likelihood of neuronal discharge and/or synaptic transmission (Monai and Hirase 2016, 2017; Monai et al. 2016). It was reported that tDCS induces cerebral plasticity based on calcium (Ca2+) elevation of astrocytic origin in mice. This tDCS-induced Ca2+ elevation is blocked by noradrenergic neuron ablation. Together, these imply that tDCS results in increased Noradrenaline (NA) release (Monai and Hirase 2016, 2017; Monai et al. 2016). The question is whether enhanced attentional orienting and inhibitory control can be explained by increased noradrenergic neurotransmission. Previous studies in psychopharmacology lend support to this notion. In a study, noradrenergic challenge with Clonidine resulted in a reduction of attentional bias, as indexed by electrophysiology. Hence, inversely, the facilitation of NA plausibly results in enhanced attentional bias, which aligns with the current observed results in the study(Logemann et al. 2014b). On the other hand, another study on inhibitory control indicated that inhibitory activity (evidenced by the Stop P3) is reduced following attenuated NA neurotransmission(Logemann et al. 2013). Therefore, tDCS enhancement of NA would likely result in improved inhibitory control simultaneously.

Lastly, the only significant behavioral effect in the experiment was the main effect of condition on attentional disengagement. We unsurprisingly found that attentional disengagement was reduced in the reward condition compared to the neutral condition regardless of group. In the absence of a significant effect from tDCS, we can expect food stimuli to pose a challenge to attentional disengagement potentially. This finding is in line with similar studies reporting that food stimuli challenge inhibitory control (Jasinska et al. 2012; Houben et al. 2014).

Despite the valuable insights gained from this study, several limitations need to be addressed. First, as indicated previously, individual variations in brain anatomy, coupled with a relatively modest sample size, might have contributed to the null results, and discrepancy between frequentist and Bayesian approaches in attentional disengagement outcomes. Besides, the effectiveness of tDCS is contingent on various factors, including stimulation parameters (e.g., duration, intensity, electrode placement). While the chosen intervention parameters were based on existing literature and rationale, other stimulation settings could have yielded different results. In the current study, the tDCS intervention was administered in a single session. Multiple sessions might produce different effects on frontal brain activity, potentially influencing the observed outcomes. Although FAA was investigated as a primary mediator, this study highlights that other neurobiological/neurochemical mechanisms, such as the noradrenergic system, may underlie the observed effects regarding attentional reorienting. However, the specific mechanisms were not directly explored in this study, leaving room for further investigation. Overall, while the study contributes necessary groundwork in this area, the identified limitations should be taken into consideration while interpreting the findings and for future research.

## Conclusion

In conclusion, this preregistered study explored the relationship between frontal alpha asymmetry (FAA), approach tendencies, and transcranial direct current stimulation (tDCS) within a visuospatial cueing paradigm. While tDCS did not show significant effects on FAA or behavioral attentional processes, intriguing secondary findings suggest tDCS might enhance cue-induced approach tendencies in a reward context. However, these effects did not lead to observable behavioral changes. Our study highlights the potential involvement of the noradrenergic system in the effect of tDCS on attentional orienting to reward-associated stimuli.

## Data Availability Statement

All data and codes can be found on our online public repository.

## References

Akil AM, Cserjési R, Nagy T, Demetrovics Z, Németh D, Logemann HNA. 2023. No Evidence of Transcranial Direct Current Stimulation on Frontal Brain Asymmetry and Inhibitory Control: A Randomized Controlled Trial. bioRxiv. 2023.09.20.558664.

Amodio DM, Master SL, Yee CM, Taylor SE. 2008. Neurocognitive components of the behavioral inhibition and activation systems: Implications for theories of self-regulation. Psychophysiology. 45:11–19.

Barry RJ, Clarke AR, Johnstone SJ, Magee CA, Rushby JA. 2007. EEG differences between eyes-closed and eyes-open resting conditions. Clin Neurophysiol. 118:2765–2773.

Baumeister RF, Heatherton TF, Tice DM. 1994. Losing control: How and why people fail at self-regulation., Losing control: How and why people fail at self-regulation. San Diego, CA, US: Academic Press.

Clark CR, Geffen GM, Geffen LB. 1989. Catecholamines and the covert orientation of attention in humans. Neuropsychologia.

Clarke P, Macleod C, Guastella A. 2011. Assessing the role of spatial engagement and disengagement of attention in anxiety-linked attentional bias: a critique of current paradigms and suggestions for future research directions. Anxiety Stress Coping. 26.

Coan JA, Allen JJB, McKnight PE. 2006. A capability model of individual differences in frontal EEG asymmetry. Biol Psychol. 72:198–207.

Common European Framework of Reference for Languages: Learning, Teaching, Assessment. 2001. Cambridge: Cambridge University Press.

Corbetta M, Patel G, Shulman GL. 2008. The reorienting system of the human brain: from environment to theory of mind. Neuron. 58:306–324.

Corbetta M, Shulman GL. 2002. Control of goal-directed and stimulus-driven attention in the brain. Nat Rev Neurosci. 3:201–215.

Faul F, Erdfelder E, Buchner A, Lang A-G. 2009. Statistical power analyses using G*Power 3.1: Tests for correlation and regression analyses. Behav Res Methods. 41:1149–1160.

Faul F, Erdfelder E, Lang A-G, Buchner A. 2007. G*Power 3: A flexible statistical power analysis program for the social, behavioral, and biomedical sciences. Behav Res Methods. 39:175–191.

Gratton G, Coles MGH, Donchin E. 1983. A new method for off-line removal of ocular artifact. Electroencephalogr Clin Neurophysiol. 55:468–484.

Harmon-Jones E, Gable PA, Peterson CK. 2010. The role of asymmetric frontal cortical activity in emotion-related phenomena: A review and update. Biol Psychol. 84:451–462.

Heatherton TF, Wagner DD. 2011. Cognitive neuroscience of self-regulation failure. Trends Cogn Sci. 15:132–139.

Houben K, Nederkoorn C, Jansen A. 2014. Eating on impulse: The relation between overweight and food-specific inhibitory control. Obesity. 22:2013–2015.

IBM Corporation. n.d. IBM SPSS Statistics for Windows.

Iyer MB, Mattu U, Grafman J, Lomarev M, Sato S, Wassermann EM. 2005. Safety and cognitive effect of frontal DC brain polarization in healthy individuals. Neurology. 64:872 LP – 875.

Jasinska AJ, Yasuda M, Burant CF, Gregor N, Khatri S, Sweet M, Falk EB. 2012. Impulsivity and inhibitory control deficits are associated with unhealthy eating in young adults. Appetite. 59:738–747.

Johnstone T, Van Reekum C, Urry H, Kalin N, Davidson R. 2007. Johnstone T, van Reekum CM, Urry HL, Kalin NH, Davidson RJ. Failure to regulate: counterproductive recruitment of top-down prefrontal-subcortical circuitry in major depression. J Neurosci 27: 8877–8884. J Neurosci. 27:8877–8884.

Kappenman E, Luck S. 2012. The Oxford Handbook of Event-Related Potential Components.

Kekic M, McClelland J, Bartholdy S, Boysen E, Musiat P, Dalton B, Tiza M, David AS, Campbell IC, Schmidt U. 2017. Single-Session Transcranial Direct Current Stimulation Temporarily Improves Symptoms, Mood, and Self-Regulatory Control in Bulimia Nervosa: A Randomised Controlled Trial. PLoS One. 12:e0167606.

Kelley NJ, Hortensius R, Schutter DJLG, Harmon-Jones E. 2017. The relationship of approach/avoidance motivation and asymmetric frontal cortical activity: A review of studies manipulating frontal asymmetry. Int J Psychophysiol. 119:19–30.

Laakso I, Tanaka S, Koyama S, De Santis V, Hirata A. 2015. Inter-subject Variability in Electric Fields of Motor Cortical tDCS. Brain Stimul. 8:906–913.

Laakso I, Tanaka S, Mikkonen M, Koyama S, Sadato N, Hirata A. 2016. Electric fields of motor and frontal tDCS in a standard brain space: A computer simulation study. Neuroimage. 137.

Lansbergen M, Böcker K, Bekker E, Kenemans J. 2007. Neural correlates of stopping and self-reported impulsivity. Clin Neurophysiol. 118:2089–2103.

Logan GD, Cowan WB, Davis KA. 1984. On the ability to inhibit simple and choice reaction time responses: A model and a method. J Exp Psychol Hum Percept Perform. 10:276–291.

Logemann HNA, Böcker K, Deschamps P, Kemner C, Kenemans J. 2013. The effect of noradrenergic attenuation by clonidine on inhibition in the stop signal task. Pharmacol Biochem Behav. 110.

Logemann HNA, Böcker KBE, Deschamps PKH, Harten PN van, Koning J, Kemner C. 2017. Haloperidol 2 mg impairs inhibition but not visuospatial attention. Psychopharmacology (Berl). 234:235–244.

Logemann HNA, Böcker KBE, Deschamps PKH, Kemner C, Kenemans JL. 2014a. The effect of the augmentation of cholinergic neurotransmission by nicotine on EEG indices of visuospatial attention. Behav Brain Res. 260:67–73.

Logemann HNA, Böcker KBE, Deschamps PKH, Kemner C, Kenemans JL. 2014b. The effect of attenuating noradrenergic neurotransmission by clonidine on brain activity measures of visuospatial attention. Hum Psychopharmacol Clin Exp. 29:46–54.

M.J.F. Robinson, A.M. Fischer, A. Ahuja ENLHM. 2016. Roles of“Wanting”and“Liking”in Motivating Behavior: Gambling, Food,and Drug Addictions. Curr Top Behav Neurosci. 27:105–136.

Mangun G, Hillyard S. 1991. Modulations of Sensory-Evoked Brain Potentials Indicate Changes in Perceptual Processing During Visual-Spatial Priming. J Exp Psychol Hum Percept Perform. 17:1057–1074.

Mathôt S, Schreij D, Theeuwes J. 2012. OpenSesame: An open-source, graphical experiment builder for the social sciences. Behav Res Methods. 44:314–324.

Meinke A, Thiel C, Fink GR. 2006. Effects of nicotine on visuo-spatial selective attention as indexed by event-related potentials. Neuroscience. 141:201–212.

Monai H, Hirase H. 2016. Astrocytic calcium activation in a mouse model of tDCS—Extended discussion. Neurogenesis. 3:e1240055.

Monai H, Hirase H. 2017. Astrocytes as a target of transcranial direct current stimulation (tDCS) to treat depression. Neurosci Res. 126.

Monai H, Ohkura M, Tanaka M, Oe Y, Konno A, Hirai H, Mikoshiba K, Itohara S, Nakai J, Iwai Y, Hirase H. 2016. Calcium imaging reveals glial involvement in transcranial direct current stimulation-induced plasticity in mouse brain. Nat Commun. 7:11100.

Nexus-32. n.d.

Opitz A, Paulus W, Will S, Antunes A, Thielscher A. 2015. Determinants of the electric field during transcranial direct current stimulation. Neuroimage. 109.

Posner MI, Snyder CR, Davidson BJ. 1980. Attention and the detection of signals. J Exp Psychol Gen. 109:160–174.

R software: a tool analysing experimental data. 2016.

Saturnino G, Antunes A, Thielscher A. 2015. On the importance of electrode parameters for shaping electric field patterns generated by tDCS. Neuroimage. 120.

Schestatsky P, Morales-Quezada L, Fregni F. 2013. Simultaneous EEG monitoring during transcranial direct current stimulation. J Vis Exp. 1–11.

Smith EE, Reznik SJ, Stewart JL, Allen JJB. 2017. Assessing and conceptualizing frontal EEG asymmetry: An updated primer on recording, processing, analyzing, and interpreting frontal alpha asymmetry. Int J Psychophysiol. 111:98–114.

Stoet G. 2010. PsyToolkit: A software package for programming psychological experiments using Linux. Behav Res Methods. 42:1096–1104.

Stoet G. 2017. PsyToolkit: A Novel Web-Based Method for Running Online Questionnaires and Reaction-Time Experiments. Teach Psychol. 44:24–31.

Thiel C, Fink G. 2008. Effects of the cholinergic agonist nicotine on reorienting of visual spatial attention and top-down attentional control. Neuroscience. 152:381–390.

Tremblay S, Lepage J, Latulipe-Loiselle A, Fregni F, Pascual-Leone A, Théoret H. 2014. The Uncertain Outcome of Prefrontal tDCS. Brain Stimul. 7.

Truong D, Magerowski G, Blackburn G, Bikson M, Alonso-Alonso M. 2013. Computational modeling of transcranial direct current stimulation (tDCS) in obesity: Impact of head fat and dose guidelines. Neuroimage (Amst). 2:759–766.

Tsegaye A, Guo C, Stoet G, Cserjési R, Kökönyei G, Logemann HNA. 2022. The relationship between reward context and inhibitory control, does it depend on BMI, maladaptive eating, and negative affect? BMC Psychol. 10:4.

Van Der Lubbe RHJ, Neggers SFW, Verleger R, Kenemans JL. 2006. Spatiotemporal overlap between brain activation related to saccade preparation and attentional orienting. Brain Res. 1072:133–152.

Wagenmakers E-J, Marsman M, Jamil T, Ly A, Verhagen J, Love J, Selker R, Gronau QF, Šmíra M, Epskamp S, Matzke D, Rouder JN, Morey RD. 2018. Bayesian inference for psychology. Part I: Theoretical advantages and practical ramifications. Psychon Bull Rev.

Wagenmakers E-J, Wetzels R, Borsboom D, van der Maas HLJ. 2011. Why psychologists must change the way they analyze their data: The case of psi: Comment on Bem (2011). J Pers Soc Psychol.

Watson P, Pearson D, Theeuwes J, Most SB, Le Pelley ME. 2020. Delayed disengagement of attention from distractors signalling reward. Cognition. 195:104125.

